# Leveraging the Adolescent Brain Cognitive Development Study to improve behavioral prediction from neuroimaging in smaller replication samples

**DOI:** 10.1101/2023.06.16.545340

**Authors:** Carolina Makowski, Timothy T. Brown, Weiqi Zhao, Donald J. Hagler, Pravesh Parekh, Hugh Garavan, Thomas E. Nichols, Terry L. Jernigan, Anders M. Dale

## Abstract

Magnetic resonance imaging (MRI) is a popular and useful non-invasive method to map patterns of brain structure and function to complex human traits. Recently published observations in multiple large scale studies cast doubt upon these prospects, particularly for prediction of cognitive traits from structural and resting state functional MRI, which seems to account for little behavioral variability. We leverage baseline data from thousands of children in the Adolescent Brain Cognitive Development^SM^ (ABCD^®^) Study to inform the replication sample size required with both univariate and multivariate methods across different imaging modalities to detect reproducible brain-behavior associations. We demonstrate that by applying multivariate methods to high-dimensional brain imaging data, we can capture lower dimensional patterns of structural and functional brain architecture that correlate robustly with cognitive phenotypes and are reproducible with only 41 individuals in the replication sample for working memory-related functional MRI, and ∼100 subjects for structural MRI. Even with 100 random re-samplings of 50 subjects in the discovery sample, prediction can be adequately powered with 98 subjects in the replication sample for multivariate prediction of cognition with working memory task functional MRI. These results point to an important role for neuroimaging in translational neurodevelopmental research and showcase how findings in large samples can inform reproducible brain-behavior associations in small sample sizes that are at the heart of many investigators’ research programs and grants.

## INTRODUCTION

Understanding how the brain gives rise to behavior is a central goal and challenge of modern neuroscience. Non-invasive neuroimaging techniques have yielded valuable opportunities to link brain structure and function with cognitive and mental health phenotypes, which in turn could be useful to predict later behavioral outcomes (1). However, recent reports have provided evidence that call into question the reproducibility of brain-behavior associations across various magnetic resonance imaging (MRI) modalities (2–6), even in large samples such as the Adolescent Brain Cognitive Development^SM^ Study (ABCD Study^®^), which have yielded much smaller effect sizes than anticipated (7).

Recent reports have suggested that studies require thousands of individuals to be sufficiently powered to measure reproducible brain-behavior associations when applying commonly used univariate statistical methods (3, 6). This has resulted in significant concerns across the brain imaging community, and beyond, about the value of MRI in trying to understand human behavior, particularly for the preponderance of investigations that have been conducted in sample sizes well under one hundred, let alone thousands, of individuals. Dozens of research groups and media outlets have independently responded to this highly cited claim of thousands of individuals being required in brain-wide association studies, including many commentaries (8–18), reports using synthetic data (19, 20), neuroimaging/cognitive data from the UK Biobank and/or Human Connectome Project (19–21), and psychopathology/genetic data from the ABCD Study (22). Across this wave of responses, it has become increasingly clear that hundreds of individuals should be sufficient to measure reproducible brain-behavior associations, particularly with the right choice of multivariate prediction methods (21) and a more intentional focus on specific behavioral phenotypes.

It remains to be seen how such multivariate methods enhance power to predict behavior in a neurodevelopmental sample such as the ABCD Study. Further, many reports focusing on brain-behavior reproducibility issues have focused on resting state functional MRI, necessitating a broader investigation of different imaging modalities, including structural and diffusion MRI, and task-evoked brain activation with fMRI. Marek, Tervo-Clemmens and colleagues (3) examined both multivariate methods and task fMRI, but did not highlight these results in the main conclusions of their paper, and used a multivariate technique that may not have been optimal in predicting cognition (21). Task fMRI (tfMRI) is of particular interest in the prediction of cognitive performance, given its success in mapping patterns of brain activity evoked by different tasks, identifying both distributed and specific neural responses to different processing demands. Our group and others have recently shown that task-evoked activation outperforms resting state functional connectivity measures when predicting cognition in the ABCD study (23–25), and similar conclusions have been drawn in other samples of youth and adults (26–29).

Here we compare the sample sizes required to detect reproducible brain-behavior associations across imaging modalities using univariate and multivariate methods. We demonstrate that by applying multivariate prediction methods that are sensitive to the complex familial structure of the ABCD study, we can capture significant patterns of associations between behavior and lower dimensional representations of structural/functional brain architecture, reducing the burden of multiple comparisons common to univariate studies. Multivariate analyses in turn yield larger effect sizes and more reproducible patterns of brain-behavior associations, negating the need for thousands of individuals, and instead showing sufficient power can be reached with well under one hundred individuals.

## RESULTS

### Participants

Participants were drawn from the baseline visit of the ABCD Study, a longitudinal neuroimaging study that tracks brain and behavioral development of ∼11,880 children starting at 9-10 years of age (30, 31). Included participants and sample sizes (Supplementary Tables 1 and 3) varied by imaging modality based on recommended inclusion flags (e.g., includes quality control criteria across modalities, and behavioral performance cut-offs for tfMRI; see Supplementary Table 4). Final sample sizes per modality were: structural MRI (sMRI), *n*= 11,174 (47.96% female), mean age in months (std) = 119.04 (7.50); diffusion MRI (dMRI), *n* = 10,200 (48.44% female), mean age in months (std) = 119.11 (7.50); tfMRI, *n* = 5673 (49.29% female), mean age in months (std) = 119.80 (7.96).

### Imaging features predicting general cognition

We used five MRI-derived features (estimated at the cortical vertex level) from three imaging modalities to predict general cognitive performance, defined by the Total Composite Score from the NIH Toolbox Cognition Battery, as described by Luciana and colleagues (32). This cognitive measure allows for direct comparison with recent studies using the same outcome variable in ABCD (3, 33). A schematic of the imaging measures is shown in Supplementary Figure 1, and more details on image processing are included in previous publications (34, 35) and in Supplementary Methods. For measures derived from sMRI, we focused on cortical surface area (SA) and cortical thickness (CT). For dMRI, we used measures derived from restriction spectrum imaging (RSI) (36, 37), including restricted normalized directional diffusion (RND) within superficial white matter, and restricted normalized isotropic diffusion (RNI) intracortically. As a supplementary analysis, we also included fractional anisotropy within superficial white matter from diffusion tensor modeling as a comparison to RND (see Supplementary Material).

For tfMRI, we chose to focus on the emotional n-back (EN-back) task, which taps into working memory and emotional regulation (35, 38, 39). Our recent work showed that task-based functional connectivity and parameter estimates from the EN-back task are more predictive of general cognition than the other two tasks within the ABCD protocol, i.e., the stop signal task and monetary incentive delay task (23). We applied standard processing of time series data, assuming a single fixed impulse response function across task conditions and brain regions. For each participant, we extracted the task model parameters derived from a general linear model (GLM) applied to the time series data, which included the beta estimates of the task condition regressors (34) for the contrast between 2-back and 0-back trials, irrespective of the type of stimulus presented (i.e. emotional face or place). An average of the task model parameters across two runs was calculated.

### Univariate associations and absolute maximum correlations

Univariate beta estimates between each imaging measure and general cognition were estimated using ordinary least squares regression at each vertex. Age, sex, scanner ID and software version were used as covariates to remain consistent with other recent work (3, 23, 33, 40). Resultant cortical maps depicting brain-behavioral correlations are in **Figure 1**. Generally, correlations between brain structure and general cognition were weak (max |*r*| = 0.151, 0.150, 0.122 for CT, RND, and RNI, respectively), with slightly larger effect sizes for SA globally with total composite scores (SA max |*r*| = 0.215).

**Figure 1.**
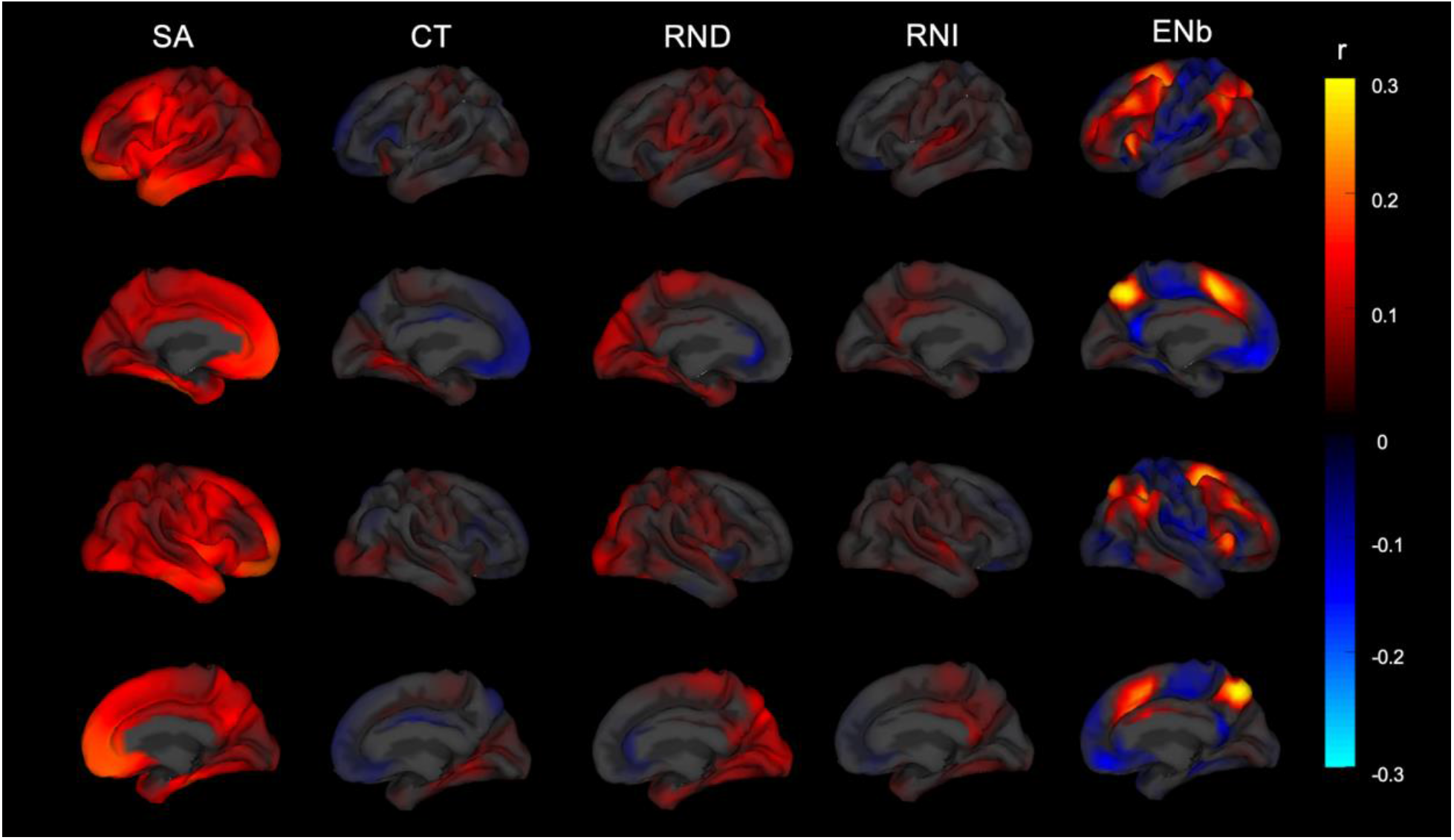
Univariate associations between five cortical features and general cognition. Abbreviations: SA. surface area; CT. cortical thickness; RND. Restricted directional diffusion within superficial white matter; RNI. Restricted isotropic diffusion intracortically; ENb. Emotional N-back, reflecting the 2-vs. 0-back contrast.

The strongest associations between the chosen imaging measures and general cognition emerged for EN-back 2-vs. 0-back task parameter estimates (EN-back max |*r*| = 0.287). Notable positive associations emerged with dorsolateral and medial prefrontal cortices and precuneus bilaterally predicting total composite scores.

### Multivariate associations

A repeated hold-out validation scheme with 100 random subsamples was used to estimate the out-of-sample prediction performance of each imaging measure on general cognition. Each input imaging measure included 5124 features (reflecting the number of vertices). For each of the 100 iterations, 90% of the sample was randomly assigned to the discovery sample and the remaining 10% to replication. Participants from the same family were kept within the same training and testing set during the cross validation. Within each training set, the mass univariate beta estimates between each imaging measure and general cognition were estimated with ordinary least squares regression, as described above for univariate analyses. Principal component analysis (PCA) and ridge regression using the Matlab function *fitrlinear* were utilized together to predict the behavioral outcome of the unseen, test-set participants. More details can be found in Supplementary Methods. The mean correlation across 100 iterations between the predicted and the observed behavioral score was used as the metric for out-of-sample behavioral prediction performance of each imaging measure and shown in **Figure 2A**.

**Figure 2.**
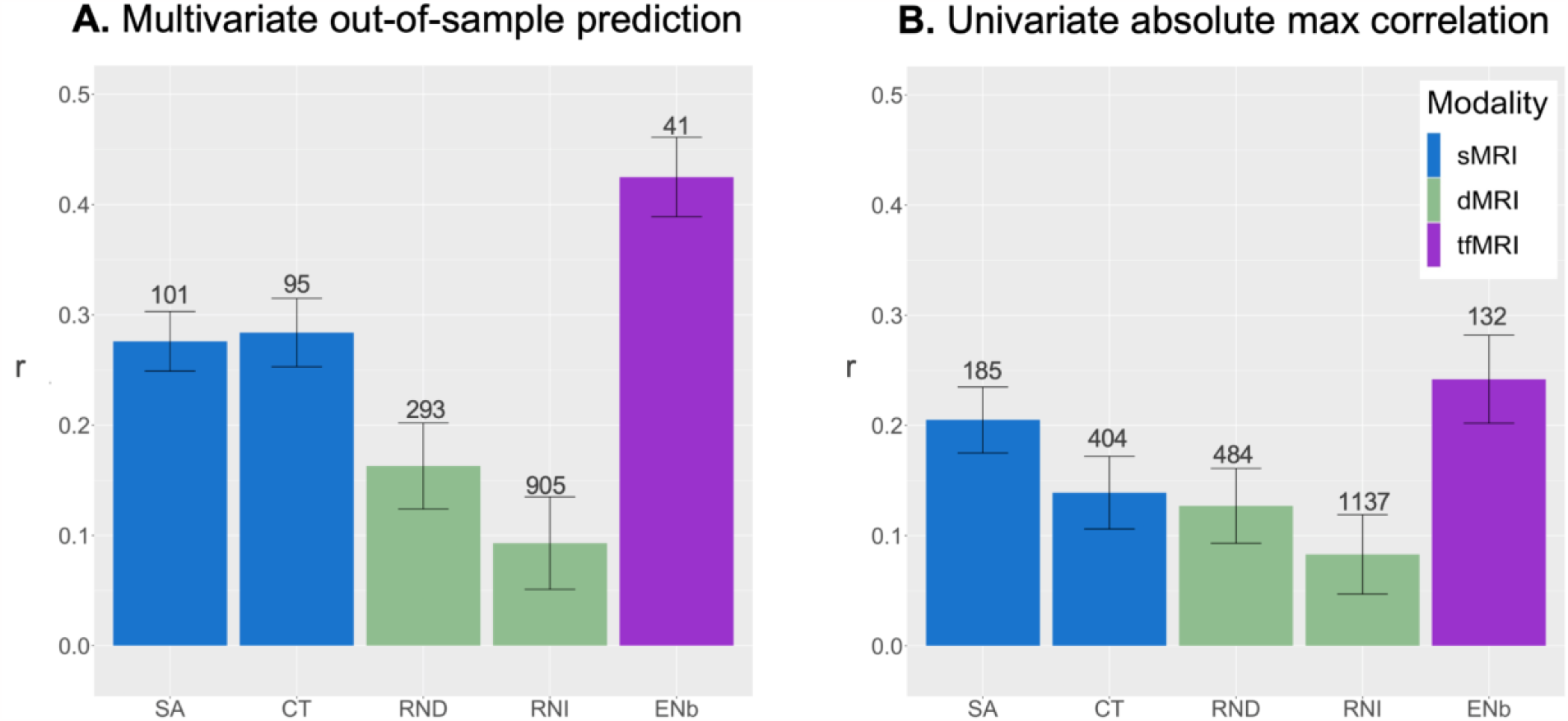
Comparison of out-of-sample prediction performance from multivariate analyses (Panel A) per imaging measure/modality, vs. maximum absolute correlations derived from univariate analyses (Panel B). Error bars reflect standard deviation, adjusted for the 10% sample overlap in test datasets. Numbers above each bar reflect the sample size required to achieve 80% power to detect effects in a replication sample, given the uncovered *r* values from the ABCD discovery sample. Abbreviations: sMRI. structural MRI; dMRI. diffusion MRI; tfMRI. task functional MRI; SA. surface area; CT. cortical thickness; RND. Restricted directional diffusion within superficial white matter; RNI. Restricted isotropic diffusion intracortically; ENb. tfMRI Emotional N-back task.

Multivariate prediction was comparable across sMRI features (SA: *r* = 0.276 ± 0.027 [adjusted standard deviation of the predicted correlation across folds, taking into consideration 10% overlap in test datasets]; CT: *r* = 0.284 ± 0.031), but still remained weak for dMRI features (RND in superficial white matter: *r* = 0.163 ± 0.039; RNI intracortically: *r* = 0.093 ± 0.042). Task fMRI estimates yielded the best prediction performance, with *r* = 0.425 ± 0.036.

For comparison, univariate correlation values were defined as the mean maximum absolute correlation value across iterations with the following outcomes (also shown in **Figure 2B**): SA: *r* = 0.205 ± 0.03; CT: *r* = 0.139 ± 0.033; RND in superficial white matter: *r* = 0.127 ± 0.034; RNI intracortically: *r* = 0.083 ± 0.036; EN-back: *r* = 0.242 ± 0.04). As expected, multivariate analyses yielded stronger effects than univariate analyses, particularly for sMRI and tfMRI measures. Only a small boost in power was observed for dMRI measures when using multivariate as compared to univariate methods, with particularly little improvement for intracortical restricted diffusion. Results were comparable for RND and FA in superficial white matter (Supplementary Figure 2), with a slightly larger boost with multivariate methods for RND (∼22% increase in prediction from univariate to multivariate) compared to FA (15% increase).

### Power calculations

Power curves for multivariate and univariate outcomes are shown in Figure 3A and 3B respectively. To achieve 80% power to detect the measured brain-behavior associations with multivariate analyses, approximately 100 subjects in the replication sample are required across sMRI features. Samples of 293 and 905 subjects are required for RND and RNI from dMRI, respectively. For EN-back tfMRI features, only 41 subjects are required. This can be compared to the higher number of subjects required with univariate associations; specifically, 185 subjects for surface area features, 404 for cortical thickness, over 480 for dMRI features, and 132 for EN-back tfMRI features. These out-of-sample replication sample sizes based on prediction performance follow classic power law principles as previously reported (21). Plots of sign error rates by sample size are included in Supplementary Figure 3.

**Figure 3.**
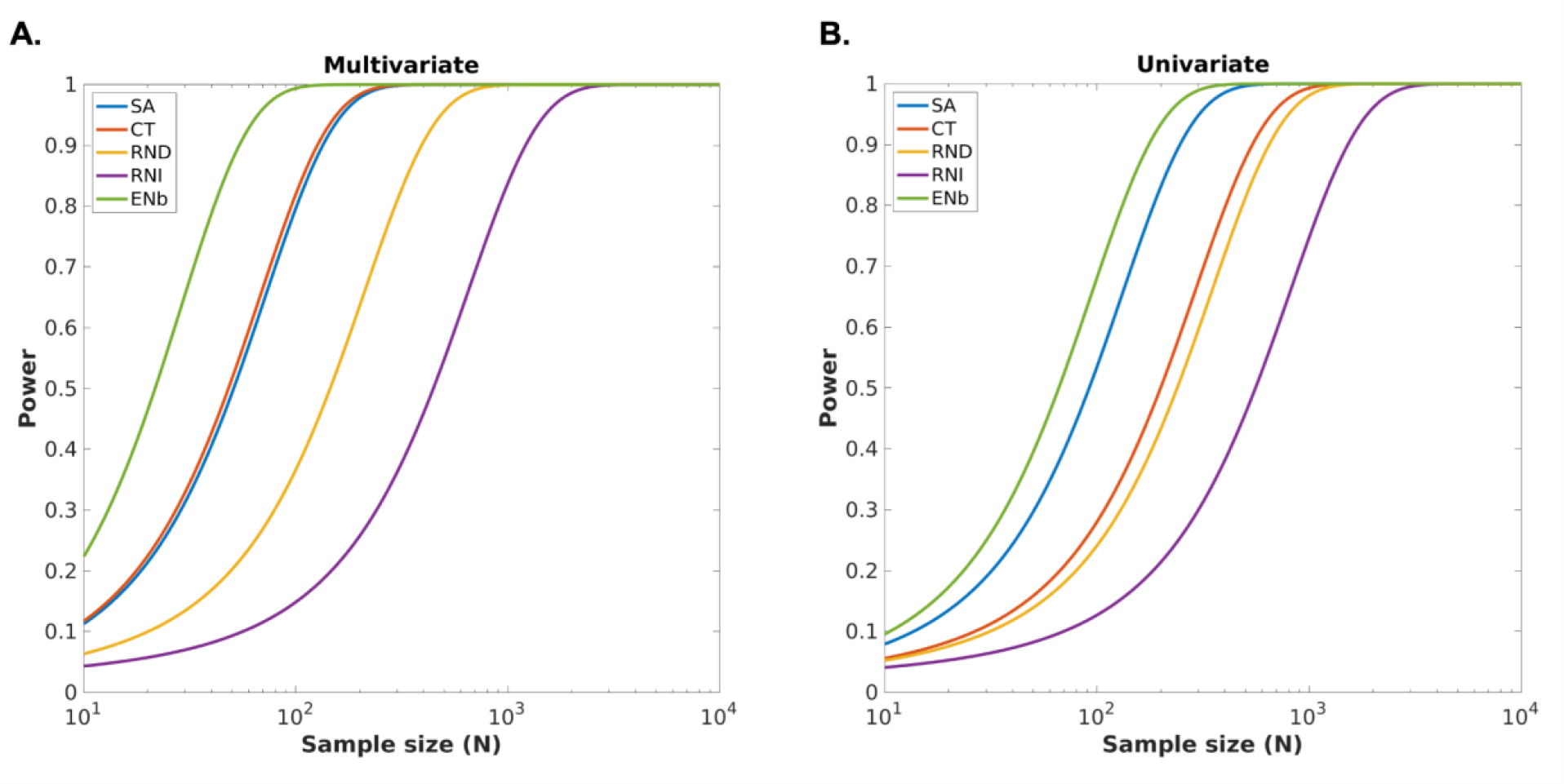
Power curves displaying replication sample sizes required (*x*-axis: log-scale N) to achieve desired level of power (*y*-axis) based on performance of each imaging measure in predicting general cognition in replication sample using (A) multivariate vs (B) univariate methods. Abbreviations: SA. surface area; CT. cortical thickness; RND. Restricted directional diffusion within superficial white matter; RNI. Restricted isotropic diffusion intracortically; ENb. tfMRI Emotional N-back task.

### Prediction performance as a function of discovery sample size

The above analyses define the replication sample size required to achieve a desired level of power, given the thousands of individuals used in the discovery sample. Our final set of analyses explored the out-of-sample prediction performance achieved over nine different discovery sample sizes spanning from 10 to 5000 participants in log units (*n* = 10, 22, 47, 103, 224, 486, 1057, 2299, 5000) for prediction of cognition from imaging features (**Figure 4**, Supplementary Figure 4), with the same prediction scheme as described above, with the remaining participants not in the discovery sample assigned to the replication sample. Consistent with the results presented above, a clear advantage for multivariate as compared to univariate methods was observed for sMRI (**Figure 4A)** and tfMRI features (**Figure 4B)**. For example, with ∼50 subjects in the discovery sample, a prediction performance of r = 0.28 was obtained for multivariate methods using ENb tfMRI features, corresponding to approximately 98 subjects in the replication sample required to achieve 80% power. This can be compared to r = 0.09 with univariate methods (∼967 subjects required) given 50 subjects in the discovery sample.

**Figure 4.**
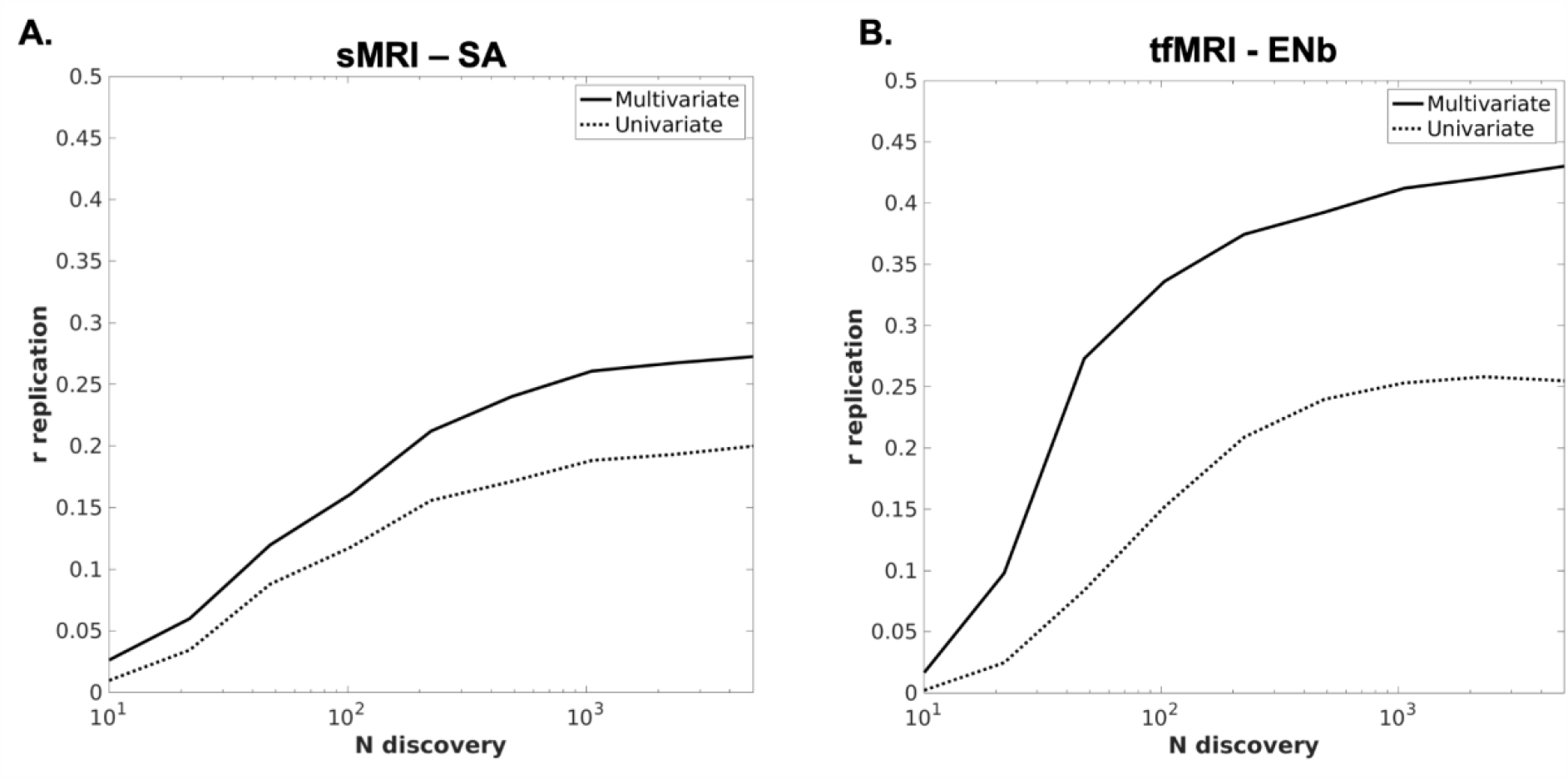
Replication curves showing out-of-sample prediction performance (defined by *r* on the *y*-axis) of A. sMRI-based surface area (SA) and B. EN-back tfMRI predicting general cognition as a function of sample size in the discovery sample (represented on the *x*-axis in log-scale units). Replication curves for the remaining three imaging measures are in Supplementary Figure 4. Multivariate metrics are compared to univariate *r*-values, reflecting the absolute maximum correlation value.

### Prediction performance for other phenotypes

The out-of-sample prediction performance for seven other measures (1 in-scanner behavioral measure of 2-back trial accuracy, 3 cognitive and 3 measures related to psychopathology; Supplementary Table 2) beyond the total composite score from the NIH toolbox are shown in Supplementary Figure 5. In general, imaging measures (especially tfMRI and structural MRI) were more predictive of cognitive variables than psychopathology, although psychopathology measures still benefit from multivariate methods, especially for prediction from EN-back tfMRI and surface area for a general *‘p’* factor of psychopathology. The highest performance was achieved with EN-back tfMRI predicting 2-back trial accuracy with an r=0.503, which would require only 29 subjects in a replication sample to detect with 80% power. This finding is consistent with our previous report of in-scanner task-relevant behavior having notably strong associations with tfMRI, as compared to an out-of-scanner cognitive task (23).

## DISCUSSION

Our work demonstrates that by leveraging large datasets such as ABCD, reproducible brain-behavior associations can be measured with multivariate methods applied to structural and functional MRI in smaller replication samples (e.g., approximately 100 subjects) at the core of many existing publications, grants, and databases. We find that a working-memory functional MRI task is particularly well-powered to predict general cognition in the baseline ABCD sample, with only ∼41 subjects required to achieve 80% power, and this is further boosted to just 29 subjects when predicting a task-relevant behavior. Finally, we show that with 100 random samplings of just 50 subjects in the discovery sample, prediction can be adequately powered with just 98 subjects in the replication sample when using multivariate methods to predict cognition from working memory task-fMRI data. This work paints a much more optimistic landscape of opportunities offered by non-invasive MRI techniques across different imaging modalities with even several dozen subjects, particularly with targeted experimental designs and improved statistical analysis.

The boost in power afforded by multivariate as compared to univariate methods in measuring brain-behavior associations is well-documented (3, 21, 33, 41, 42). Associations between cognitive performance and neuroimaging phenotypes are particularly amenable to multivariate methods, given the distributed and sparse nature of these effects across the brain (41, 42). Functional MRI analysis has long capitalized on observed strong patterns of spatial and temporal correlations across the brain (43). Structural imaging-derived phenotypes also have a strong covariance structure between regions (44), likely due to gradients of core neurodevelopmental genetic factors shaping the cortex early in life (45–48). In line with our previous work (42), multivariate methods capturing distributed brain structural patterns explained a larger amount of variance in cognition compared to univariate mapping. We observed weak univariate effect sizes between structural/diffusion imaging measures and cognitive measures, with the lowest effects for intracortical restricted diffusion. However, and importantly, these small effects with sMRI received substantial boosts with multivariate modeling. Our results emphasize the importance of taking into consideration structural brain patterns as a whole in brain-behavior analyses, rather than analyzing any single vertex or region-of-interest in isolation. Although results were weaker overall with measures of psychopathology (e.g., internalizing and externalizing symptoms, *p* factor of psychopathology), these measures still benefited from a boost in prediction, particularly when predicted by tfMRI and surface area, using multivariate methods. Generally, we also observed that surface area and task fMRI performed quite well for other cognitive measures, once again suggesting that replication sample sizes still only require on the order of hundreds, not thousands, of individuals for these imaging measures.

We did not observe large boosts in effect sizes with diffusion MRI measures when comparing multivariate to univariate associations. RND within superficial white matter showed a slight boost, with 290 participants required in the replication sample to achieve 80% power in predicting cognition with multivariate methods, compared to 508 with univariate methods at the same level of power. Notably, RND, derived from restriction spectrum imaging modeling, benefited more from multivariate methods for prediction of general cognition compared to fractional anisotropy of superficial white matter. The slight improvement in performance of RND compared to FA is consistent with several reports that RSI measures, derived from multi-shell diffusion imaging acquisitions, may be more sensitive in capturing microstructural properties especially in the context of neurodevelopment (36), and neurological and psychiatric disorders (49–51). For restricted isotropic diffusion intracortically, both methods still required over 1000 individuals. Although the current manuscript focused on peri-cortical associations (i.e. in and around the cortex), it cannot be ruled out that diffusion measures of other brain regions, that may be more reliably measured with MRI (e.g., deeper white matter tracts, subcortical structures), would still benefit from multivariate analyses. Future work extending multivariate analyses to voxel-wise prediction of cognitive and other behavioral phenotypes hypothesized to be more strongly associated with microstructure would be fruitful in testing this hypothesis.

The conclusion reaching many headlines that thousands of individuals are required to measure brain-behavior associations with structural and functional MRI (3) is based on the assumption that researchers are working within a univariate framework and sampling across a broad array of outcomes. This conclusion was also based largely on the poor performance of resting-state functional connectivity to predict cognition and other behaviors, a method that does not orient participants’ attention to any particular task or stimulus, and in turn, may not be the best candidate for predicting measures of cognition (9, 23). This ‘brain-wide association study’ approach of sampling across a large array of behavioral outcomes with mass univariate analyses are not unlike the methods used in genome-wide association studies (GWAS). However, the application of multivariate methods helps ease the burden of multiple comparison corrections that mass univariate methods typically require. Instead, our findings highlight a different use case of a GWAS-like framework, where we can leverage a larger discovery sample, such as the one afforded by the ABCD Study, to pinpoint multivariate brain patterns associated with behavioral measures of interest, and in turn, inform findings in smaller samples. In this vein, our approach parallels that of the application of consortia-led GWAS to the derivation of polygenic risk scores in independent datasets, where large samples can help us obtain more precise predictive weights for out-of-sample prediction.

However, large ‘ABCD-like’ samples are not always necessary if working with strong brain-behavior relationships to begin with. Here we highlight analyses testing out-of-sample prediction performance given different discovery sample sizes, showing that even with 100 random samplings of 50 subjects in the discovery sample, only ∼100 subjects are required in replication to predict general cognition from task fMRI data using multivariate methods. Additionally, integration of regularization methods in our multivariate prediction regime may have contributed to the well-powered results even in smaller discovery samples by reducing the chances of overfitting, leading to a more optimal model. This further emphasizes the fact that with the correct methodological choices, thousands of individuals are not always required in either discovery or replication samples in pursuit of meaningful and reproducible brain-behavior associations. This is particularly evident when predicting relevant in-scanner behavior (i.e. 2-back accuracy) from task activation during the EN-back task, where only 29 replication subjects were required to assess this relationship with 80% power. Larger samples, however, are required for other weaker brain-behavior associations, for instance with internalizing and externalizing measures of psychopathology in ABCD.

We recognize that although effects were strongest with task fMRI compared to other imaging modalities, the included sample of 5673 individuals may not generalize to the full sociodemographic spread of the ABCD baseline sample. We aimed to stay as consistent as possible with other recent work testing the reproducibility and power of brain-behavior associations with neuroimaging, and restricted our covariates to age, sex, and scanner type. However, future investigations would benefit from a deeper understanding of the impact of sociodemographic and environmental factors in behavioral prediction paradigms. We also note that our task fMRI sample retained approximately double the number of participants compared to published investigations using resting state fMRI in ABCD (3, 23, 33). The notable drop in subjects with resting-state fMRI may be in part due to its experimental design, where participants do not need to engage in a goal-oriented task and may be more likely to move in the scanner, thus limiting its generalizability to the broader ABCD sample (52). Given these limitations and relatively weak predictive power of resting-state fMRI compared to task fMRI for cognition (3, 41), there has been increased dialogue surrounding a potential paradigm shift from resting state to task-based functional MRI measures (27). Mental health-related phenotypes are of particular interest in adolescent samples, but show weaker effects with brain structure and function in ABCD Study data compared to general cognition, both in this work and from others (3). Future directions would benefit from a focus on modeling of behavioral phenotypes to improve precision behavioral phenotyping (22). This work focused on cross-sectional baseline data, but future work is encouraged to integrate longitudinal data from ABCD to boost effect sizes of brain-behavior associations (53), as well as integrate independent datasets for external validation. It should also be noted that the ABCD Study integrates a complex study design, including combining data across 21 different sites. Thus, prediction performance in such large multi-site studies may actually be underestimating true brain-behavior effects that could be more precisely studied in single-site/scanner studies that many grants are based on. Finally, recommendations have been made to steer away from single modeling approaches and understand how various models (e.g. mixture modeling) (54) could better capture the complex relationships between brain and behavior reflected in large diverse datasets such as ABCD.

We show that reproducible brain-behavior associations can be obtained with dozens, rather than thousands, of individuals, helping to quell growing concerns of the demise of reproducible results with neuroimaging. Our findings help shed clarity on the utility of neuroimaging studies, particularly for understanding normative and aberrant neurodevelopment, and present a more hopeful view for future funding priorities of smaller neuroimaging studies and policy decisions.

## ACKNOWLEDGMENTS/FUNDING

The authors would like to thank the research participants and staff involved in data collection of the Adolescent Brain Cognitive Development (ABCD) Study data. The ABCD Study is a multisite, longitudinal study designed to recruit more than 10,000 children ages 9 and 10 and follow them over 10 years into early adulthood. The ABCD Study is supported by the National Institutes of Health (NIH) and additional federal partners under award numbers U01DA041048, U01DA050989, U01DA051016, U01DA041022, U01DA051018, U01DA051037, U01DA050987, U01DA041174, U01DA041106, U01DA041117, U01DA041028, U01DA041134, U01DA050988, U01DA051039, U01DA041156, U01DA041025, U01DA041120, U01DA051038, U01DA041148, U01DA041093, U01DA041089, U24DA041123, and U24DA041147. A full list of supporters is available at https://abcdstudy.org/federal-partners.html. A listing of participating sites and a complete listing of the study investigators can be found at https://abcdstudy.org/study-sites/. ABCD consortium investigators designed and implemented the study and/or provided data but did not necessarily all participate in analysis or writing of this report. This manuscript reflects the views of the authors and may not reflect the opinions or views of the NIH or ABCD consortium investigators. The ABCD data repository grows and changes over time. Data were drawn from the NIMH Data Archive ABCD Collection Release 4.0 (DOI: 10.15154/1523041).

Additionally, C.M. is supported by the National Institutes of Mental Health, award number 1K99MH132886. P.P. is supported by funding from the European Union’s Horizon 2020 research and innovation programme under the Marie Skłodowska-Curie grant agreement number 801133 and from the Research Council of Norway grant number 324252.

## REFERENCES

1. O. Kardan, et al., Differences in the functional brain architecture of sustained attention and working memory in youth and adults. PLOS Biology 20, e3001938 (2022).

2. M. L. Elliott, et al., What Is the Test-Retest Reliability of Common Task-Functional MRI Measures? New Empirical Evidence and a Meta-Analysis. Psychol. Sci. 31, 792–806 (2020).

3. S. Marek, et al., Reproducible brain-wide association studies require thousands of individuals. Nature 603, 654–660 (2022).

4. J. T. Kennedy, et al., Reliability and stability challenges in ABCD task fMRI data. Neuroimage 252, 119046 (2022).

5. R. E. Kelly Jr, M. J. Hoptman, Replicability in Brain Imaging. Brain Sci 12 (2022).

6. S. Liu, A. Abdellaoui, K. J. H. Verweij, G. van Wingen, Replicable brain-phenotype associations require large-scale neuroimaging data. Nat Hum Behav 7, 1344–1356 (2023).

7. A. S. Dick, et al., Meaningful associations in the adolescent brain cognitive development study. Neuroimage 239, 118262 (2021).

8. X.-Z. Kong, C. Zhang, Y. Liu, Y. Pu, Scanning reproducible brain-wide associations: sample size is all you need? Psychoradiology 2, 67–68 (2022).

9. M. D. Rosenberg, E. S. Finn, How to establish robust brain–behavior relationships without thousands of individuals. Nat. Neurosci. 25, 835–837 (2022).

10. Revisiting doubt in neuroimaging research. Nat. Neurosci. 25, 833–834 (2022).

11. C. Gratton, S. M. Nelson, E. M. Gordon, Brain-behavior correlations: Two paths toward reliability. Neuron 110, 1446–1449 (2022).

12. C. G. Deyoung, et al., Reproducible between-person brain-behavior associations do not always require thousands of individuals. PsyArXiv (2022) 10.31234/osf.io/sfnmk (April 4, 2023).

13. M. Mallar Chakravarty, Controversies on brain-wide association studies: commentaries from the field. Aperture Neuro, 2 (2022).

14. Cognitive neuroscience at the crossroads. Nature 608, 647 (2022).

15. P. A. Bandettini, et al., The challenge of BWAs: Unknown unknowns in feature space and variance. Med 3, 526–531 (2022).

16. J. Tiego, A. Fornito, Putting behaviour back into brain-behaviour correlation analyses Aperture Neuro, BWAS Editorials, 1–4 (2022).

17. L. Q. Uddin, Brain–behavior associations depend heavily on user-defined criteria. Aperture Neuro, 2 (2022).

18. S. L. Valk, M. Hettwer, Commentary on ‘Reproducible brain-wide association studies require thousands of individuals’ Aperture Neuro, 2 (2022).

19. L. Cecchetti, G. Handjaras, Reproducible brain-wide association studies do not necessarily require thousands of individuals. PsyArXiv (2022).

20. M. Gell, et al., The Burden of Reliability: How Measurement Noise Limits Brain-Behaviour Predictions. bioRxiv (2023).

21. T. Spisak, U. Bingel, T. D. Wager, Multivariate BWAS can be replicable with moderate sample sizes. Nature 615, E4–E7 (2023).

22. J. Tiego, et al., Precision behavioral phenotyping as a strategy for uncovering the biological correlates of psychopathology. Nat Ment Health 1, 304–315 (2023).

23. W. Zhao, et al., Task fMRI paradigms may capture more behaviorally relevant information than resting-state functional connectivity. Neuroimage 270, 119946 (2023).

24. A. Omidvarnia, et al., Is resting state fMRI better than individual characteristics at predicting cognition? bioRxiv (2023).

25. J. Chen, et al., Shared and unique brain network features predict cognitive, personality, and mental health scores in the ABCD study. Nat. Commun. 13, 2217 (2022).

26. M. D. Rosenberg, et al., A neuromarker of sustained attention from whole-brain functional connectivity. Nat. Neurosci. 19, 165–171 (2016).

27. A. S. Greene, S. Gao, D. Scheinost, R. T. Constable, Task-induced brain state manipulation improves prediction of individual traits. Nat. Commun. 9, 2807 (2018).

28. E. S. Finn, P. A. Bandettini, Movie-watching outperforms rest for functional connectivity-based prediction of behavior. Neuroimage 235, 117963 (2021).

29. R. Jiang, et al., Task-induced brain connectivity promotes the detection of individual differences in brain-behavior relationships. Neuroimage 207, 116370 (2020).

30. H. Garavan, et al., Recruiting the ABCD sample: Design considerations and procedures. Dev. Cogn. Neurosci. 32, 16–22 (2018).

31. N. D. Volkow, et al., The conception of the ABCD study: From substance use to a broad NIH collaboration. Dev. Cogn. Neurosci. 32, 4–7 (2018).

32. M. Luciana, et al., Adolescent neurocognitive development and impacts of substance use: Overview of the adolescent brain cognitive development (ABCD) baseline neurocognition battery. Dev. Cogn. Neurosci. 32, 67–79 (2018).

33. C. Sripada, et al., Prediction of neurocognition in youth from resting state fMRI. Mol. Psychiatry 25, 3413–3421 (2020).

34. D. J. Hagler Jr, et al., Image processing and analysis methods for the Adolescent Brain Cognitive Development Study. Neuroimage 202, 116091 (2019).

35. B. J. Casey, et al., The Adolescent Brain Cognitive Development (ABCD) study: Imaging acquisition across 21 sites. Dev. Cogn. Neurosci. 32, 43–54 (2018).

36. C. E. Palmer, et al., Microstructural development from 9 to 14 years: Evidence from the ABCD Study. Dev. Cogn. Neurosci. 53, 101044 (2022).

37. N. S. White, T. B. Leergaard, H. D’Arceuil, J. G. Bjaalie, A. M. Dale, Probing tissue microstructure with restriction spectrum imaging: Histological and theoretical validation. Hum. Brain Mapp. 34, 327–346 (2013).

38. D. M. Barch, et al., Function in the human connectome: task-fMRI and individual differences in behavior. Neuroimage 80, 169–189 (2013).

39. A. O. Cohen, et al., When Is an Adolescent an Adult? Assessing Cognitive Control in Emotional and Nonemotional Contexts. Psychol. Sci. 27, 549–562 (2016).

40. B. Chaarani, et al., Baseline brain function in the preadolescents of the ABCD Study. Nat. Neurosci. 24, 1176–1186 (2021).

41. W. Zhao, et al., Individual Differences in Cognitive Performance Are Better Predicted by Global Rather Than Localized BOLD Activity Patterns Across the Cortex. Cereb. Cortex 31, 1478–1488 (2021).

42. C. E. Palmer, et al., Distinct Regionalization Patterns of Cortical Morphology are Associated with Cognitive Performance Across Different Domains. Cereb. Cortex 31, 3856–3871 (2021).

43. G. Derado, F. D. Bowman, C. D. Kilts, Modeling the spatial and temporal dependence in FMRI data. Biometrics 66, 949–957 (2010).

44. J. P. Lerch, et al., Mapping anatomical correlations across cerebral cortex (MACACC) using cortical thickness from MRI. Neuroimage 31, 993–1003 (2006).

45. P. Rakic, Specification of cerebral cortical areas. Science 241, 170–176 (1988).

46. P. Rakic, Evolution of the neocortex: a perspective from developmental biology. Nat. Rev. Neurosci. 10, 724–735 (2009).

47. D. van der Meer, et al., Understanding the genetic determinants of the brain with MOSTest. Nat. Commun. 11, 3512 (2020).

48. C. Makowski, et al., Discovery of genomic loci of the human cerebral cortex using genetically informed brain atlases. Science 375, 522–528 (2022).

49. E. T. Reas, et al., Microstructural brain changes track cognitive decline in mild cognitive impairment. Neuroimage Clin 20, 883–891 (2018).

50. R. Q. Loi, et al., Restriction spectrum imaging reveals decreased neurite density in patients with temporal lobe epilepsy. Epilepsia 57, 1897–1906 (2016).

51. R. A. Carper, J. M. Treiber, N. S. White, J. S. Kohli, R.-A. Müller, Restriction Spectrum Imaging As a Potential Measure of Cortical Neurite Density in Autism. Front. Neurosci. 10, 610 (2016).

52. K. T. Cosgrove, et al., Limits to the generalizability of resting-state functional magnetic resonance imaging studies of youth: An examination of ABCD Study® baseline data. Brain Imaging Behav. 16, 1919–1925 (2022).

53. K. Kang, et al., Study design features that improve effect sizes in cross-sectional and longitudinal brain-wide association studies. bioRxiv (2023).

54. A. S. Greene, et al., Brain–phenotype models fail for individuals who defy sample stereotypes. Nature 609, 109–118 (2022).

