## Supplementary for "Leveraging the Adolescent Brain Cognitive Development Study to improve behavioral prediction from neuroimaging in smaller replication samples"

**SUPPLEMENTARY INFORMATION.**

*Participants*.

Participants were drawn from the baseline visit of the ABCD Study, a longitudinal neuroimaging study that tracks brain and behavioral development of ~11,880 children starting at 9-10 years of age. The ABCD Study represents a demographically and ethnically diverse cohort of youth in the United States, and includes an embedded twin cohort and siblings. Informed consent was obtained from parents/caretakers and assent was obtained from the children. More detailed descriptions of recruitment and data collection within the ABCD Sample can be found in (1, 2). See Supplementary Tables 1 and 3 for sample details per imaging modality.

*Cognitive outcomes of interest*.

We focused on prediction of a single measure of general cognitive performance, the Total Composite Score from the NIH Toolbox Cognition Battery. The Total Composite Score is an arithmetic average of the 7 subtests from the NIH Toolbox, which includes measures of vocabulary size (Picture Vocabulary Task), single word reading ability (Oral Reading Task), rapid visual processing (Pattern Comparison Processing Speed Test), working memory capacity (List Sorting Working Memory Test), episodic memory (Picture Sequence Memory Test), attention and inhibitory control (Flanker Task), and cognitive flexibility (Dimensional Change Card Sort Task). This measure has been used in many studies across various ages (3), allowing for generalizability of our findings to many other investigations. This measure also has good test-retest reliability and validity across both children and adults (4, 5). This measure was also used as the primary outcome of interest in similarly motivated papers (6). We also explored prediction of 7 other phenotypes (Supplementary Table 2; Supplementary Figure 5), including 1 in-scanner behavioral task (accuracy on 2-back trials within the EN-back task), 3 out-of-scanner cognitive measures (crystallized [comprised of Picture Vocabulary and Oral Reading tasks] and fluid composite [comprised of Pattern Comparison Processing Speed, List Sorting Working Memory, Picture Sequence Memory, Flanker, and Dimensional Change Card Sort tasks] scores from the NIH Toolbox, and Matrix Reasoning from the Wechsler Intelligence Test for Children-V (WISC-V) (7) and 3 measures of psychopathology (internalizing and externalizing symptoms from the Child Behavior Checklist, as well as a *p* factor calculated from the Child Behavior Checklist subscale scores, as presented in Clark et al. (8)).

*MRI processing and measures.*

The ABCD MRI data were collected across 21 research sites using GE 750, Siemens Prisma, and Philips Achieva and Ingenia 3T scanners. Scanning protocols were harmonized across sites. Please see Casey et al. (9) and Hagler et al. (10) for full details of imaging acquisition and preprocessing protocols, with summaries for each modality provided below.

We focus on five cortical measures across three imaging modalities in the main manuscript, depicted in Supplementary Figure 1. All of these measures were estimated at the cortical vertex level and relied on collection of T1-weighted (T1w) structural MRI images (1mm isotropic), which were acquired with a 3D T1w inversion prepared RF-spoiled gradient echo scan. Individuals were included in analyses if they had complete data for NIH toolbox total composite scores, covariates, and met modality-specific inclusion criteria, which is combined into a single variable per modality within the *abcd_imgincl01* table from released tabulated data (i.e., sMRI: *imgincl_t1w_include* = 1; dMRI: *imgincl_dmri_include* = 1; tfmri: *imgincl_nback_include* = 1). See Supplementary Table 4 for the full list of inclusion/exclusion criteria comprising each of these modality-specific inclusion flags. These filters result in $n = 11{,}174$, $n = 10{,}200$, and $n = 7{,}793$ for sMRI, dMRI, and tfMRI, respectively. An additional 2,120 participants were dropped from tfMRI analyses due to incomplete field-of-view acquisitions and/or not having complete acquisitions across two runs.

*Structural MRI (sMRI) processing and derived measures.*
For measures derived from structural MRI, we focused on cortical surface area and thickness (Supplementary Figure 1A). Cortical surfaces were constructed from T1w structural images for each subject and segmented to calculate measures of cortical thickness and surface area along 5124 vertices using FreeSurfer v7.1.1 (11–15). Cortical maps were smoothed using a Gaussian kernel of 20 mm full-width half maximum (FWHM) and mapped into standardized spherical atlas space.

*Diffusion MRI (dMRI) processing and derived measures.*
For measures derived from diffusion MRI, we focused on fractional anisotropy (FA) within superficial white matter and intracortical restricted isotropic diffusion from restriction spectrum imaging. Restricted directional diffusion (RND) within superficial white matter and intracortical restricted isotropic diffusion (RNI) was derived from restriction spectrum imaging (RSI) modeling; details of this approach can be found in previous publications (16–18). Briefly, RSI takes advantage of a multi-shell diffusion acquisition to estimate the contribution of diffusion signal from separable pools of water within a tissue, which includes free water (e.g., CSF), hindered diffusion (e.g., mostly extracellular space), and restricted diffusion (e.g., mostly intracellular space). Of interest to this study, restricted diffusion describes water within intracellular spaces confined by cell membranes with a non-Gaussian pattern of displacement. Spherical deconvolution (SD) is used to reconstruct the fiber orientation distribution (FOD) in each voxel from the restricted compartment, where the restricted tissue compartment is modeled as a fourth order spherical harmonic (SH) function. The restricted directional measure, RND, is the norm of the SH coefficients for the second and fourth order SH coefficients (divided by the norm of the coefficients across restricted, hindered, and free water compartments). In other words, RND models diffusion emanating from multiple directions within a voxel. Our second RSI measure, RNI, refers to the spherical mean of the FOD across all orientations. Both

superficial white matter RND and intracortical RNI were sampled with linear interpolation perpendicular to the gray-white matter boundary in either direction at 0.8-2 mm depths in 0.2 mm intervals; specifically, into the white matter for FA, and into the gray matter for RNI (10). Similar to structural MRI data, cortical maps were smoothed using a Gaussian kernel of 20 mm FWHM, mapped into standardized spherical atlas space, and concatenated into matrices before being used in statistical analysis.

As a comparison to RND within superficial white matter, we also included fractional anisotropy (FA), derived from the diffusion tensor model (19, 20). Diffusion tensor parameters were calculated using a standard, linear estimation approach with log-transformed diffusion-weighted (DW) signals (20). Tensor matrices were diagonalized using singular value decomposition, obtaining three eigenvectors and three corresponding eigenvalues, from which FA could be calculated (19).

*Functional MRI (fMRI) processing and derived measures.*

*EN-back task.*

For task-fMRI, we chose to focus on the 2- vs. 0-back contrast from the emotional *n*-back task (Supplementary Figure 1B) (9, 21, 22) irrespective of the type of stimulus (i.e. emotional face, place) presented. To engage working memory, the task includes 0-back and 2-back conditions, presented in a block design. Participants were instructed to indicate with a button press whether the current stimulus matched the stimulus presented 2 trials back (2-back condition) or matched the target stimulus presented at the beginning of the block (0-back condition). Although we did not focus on emotional regulation in this study, the task stimuli included happy faces, fearful faces, neutral faces, and places, presented serially. Our recent work showed that task-based functional connectivity and parameter estimates from the EN-back task are more predictive of general cognition than the other two tasks within the ABCD protocol, i.e., the stop signal task and monetary incentive delay task; thus, we focused on the EN-back task only within this manuscript.

*Task fMRI processing.*

Task fMRI acquisitions were collected with multiband EPI with slice acceleration factor 6 (2.4 mm isotropic, TR = 800 ms). Preprocessing steps are outlined in Hagler et al (10) and Zhao et al (23). Briefly, preprocessing steps included (i) head motion correction, (ii) B0 distortion correction, (iii) gradient warping correction, (iv) within-scan motion correction, and (v) registration to T1w structural images. Initial frames (Siemens and Philips scanners: 8 TRs; GE750 DV25: 5 TRs; GE750 DV26: 16 TRs) were removed from the preprocessed task fMRI time course. Motion estimates were filtered to remove the effect of respiratory signals (24). The preprocessed time courses were normalized and sampled onto the cortical surface for each participant. Task functional MRI effects were estimated at the participant level using a general linear model (GLM) that included the stimulus timing for each task condition (10) and the temporal derivative to capture any task related changes in the fMRI time course that is not captured by our task model. The GLM modeled each task condition with a double gamma function and its first temporal derivative along with 4 nuisance regressors for baseline shifts and cubic trends and 12 regressors for the six motion estimates and their temporal derivatives. For the GLM estimation, time points with framewise displacement (FD) greater than 0.9 mm were censored (25). For our tfMRI features of interest, we used the task model parameters (i.e. the beta estimates of the task condition regressors) modeling the 2-back trials vs. 0-back trials contrast, irrespective of face/place stimulus type. Parameter estimates across run 1 and run 2 were averaged.

*Statistical Analysis.*

*Estimation of univariate associations.* Mass univariate beta estimates between each imaging measure and general cognition were calculated with ordinary least squares regression at each vertex, adjusting for the effects of nested family structure in the ABCD data and specified covariates. Age, sex, scanner ID and software version were used as covariates to remain consistent with other recent work (6, 26, 27).

*Multivariate prediction with PCA and ridge regression.*
A repeated hold-out validation scheme with 100 random subsamples was used to estimate the out-of-sample prediction performance of each imaging measure on general cognition. For each of the 100 iterations, 90% of the sample was randomly assigned to the discovery sample and the remaining 10% to replication. Participants from the same family were kept within the same training and testing set during the cross validation. Confounding effects of the same covariates used in univariate analyses (age, sex, scanner ID and software version) were removed before out-of-sample prediction. Principal component analysis (PCA) and ridge regression implemented in Matlab's *fitrlinear* was utilized to predict the behavioral outcome of the unseen, test-set participants. Normalization of imaging data before dimensionality reduction was done within the cross-validation framework. The shrinkage parameter value, $\lambda$, was set to 1, consistent with previous work(27). The $k$ value, representing the fraction of principal components used in prediction, was empirically determined as follows. First, 15 different values of $k$ spanning 0 to 1 (i.e., 0.001, 0.0016, 0.0027, 0.0044, 0.0072, 0.012, 0.019, 0.032, 0.052, 0.085, 0.14, 0.23, 0.37, 0.61, 1.00) were tested in a first round of 100 random subsamples, split into 90% and 10% discovery and replication datasets, respectively. The $k$ value that yielded the optimal prediction performance was then subsequently used in a second iteration of 100 random subsamples, from which prediction performance was evaluated. The empirically determined $k$ value per imaging modality and behavioral measure is listed in Supplementary Table 3. The correlation between the predicted and the observed behavioral score was used as the metric for out-of-sample behavioral

prediction performance of each imaging measure. The standard deviation of the correlation estimates took into consideration the expected 10% overlap in replication datasets given the 90/10 split utilized in our cross-validation scheme. Specifically, all standard deviation estimates were scaled up by a factor of ~1.05, based on the finite population correction $1/\sqrt{(1-p)}$ (where *p* is the expected proportion overlap between replication samples of 0.1) (28).

*Generation of power curves*. Power curves were generated in Matlab, where the replication sample size needed to achieve a desired level of power was defined as follows (29) and consistent with recent papers (6, 27):

$$N = [(Z_\alpha+Z_\beta)/C]^2 + 3$$

where $Z_\alpha$ represents the standard normal score for a given two-tailed alpha level (set to 0.05 in our work); $Z_\beta$ represents the standard normal score for a given beta-value (e.g., 0.2 for 80% power) and C=arctan(r), where *r* represents the absolute maximum correlation for univariate analyses, taken as the mean correlation across 100 iterations of out-of-sample prediction for multivariate analyses.

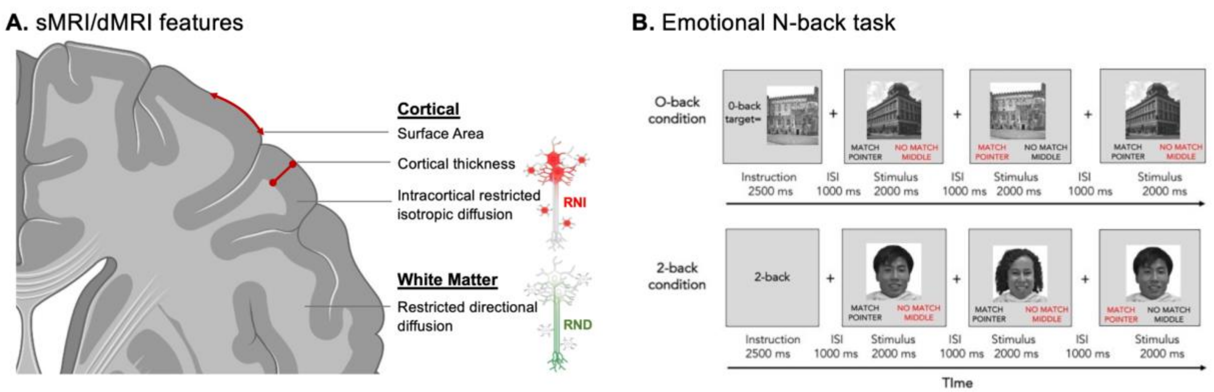


**Supplementary Figure 1**. Imaging-derived measures from A. Structural and diffusion MRI; B. task fMRI using the Emotional N-back task. Task fMRI parameter estimates were obtained from the 2- vs. 0-back contrast, and averaged across two runs, assuming a single fixed impulse response function across task conditions and brain regions.

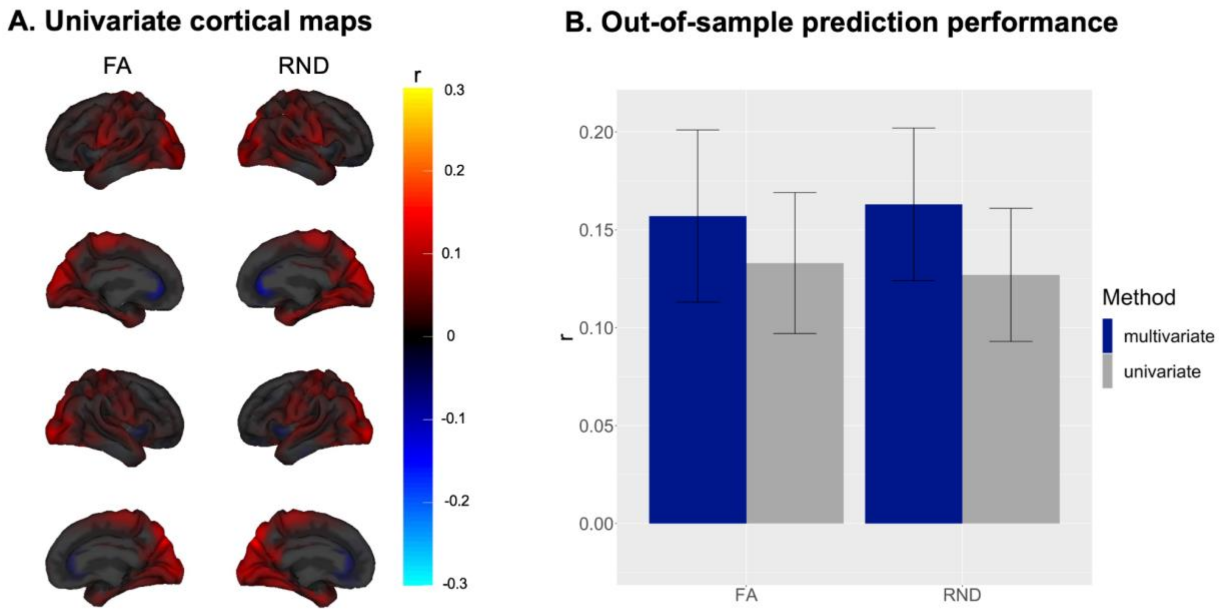


**Supplementary Figure 2.** Similar results are obtained for fractional anisotropy (FA) and restricted normalized directional diffusion (RND) measured within superficial white matter, as shown by A) univariate associations between each brain phenotype, measured within superficial white matter, and general cognition; and B) multivariate and univariate prediction of cognition. A slightly larger boost is achieved with multivariate prediction of cognition from RND (~22% increase) compared to FA (~15% increase).

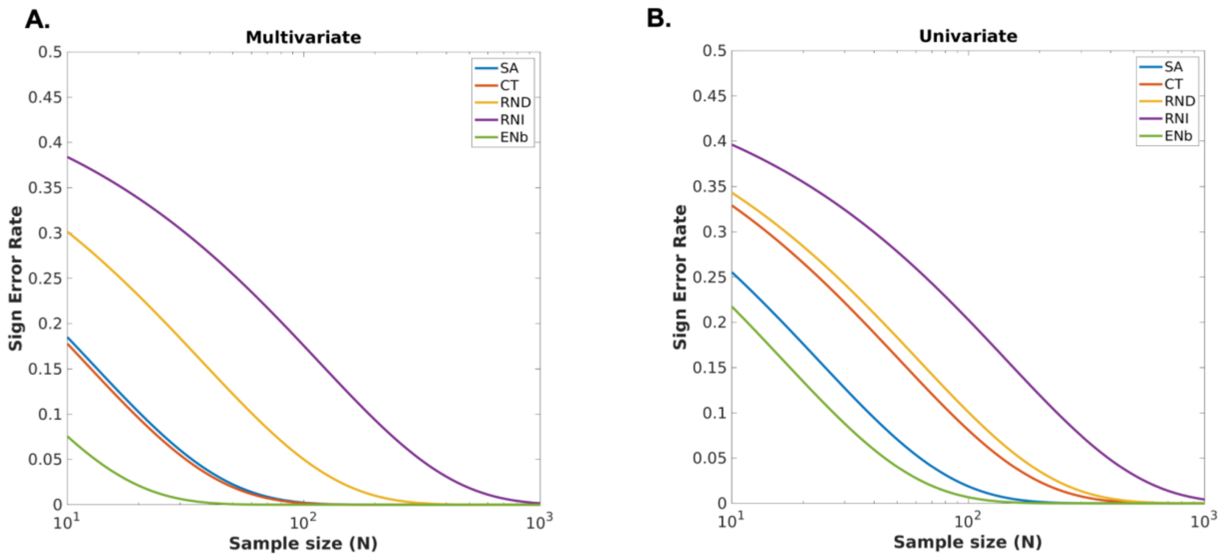


**Supplementary Figure 3**. Sign error curves (proportion of correlations with opposite sign for a given sample size) for A) multivariate and B) univariate analyses.

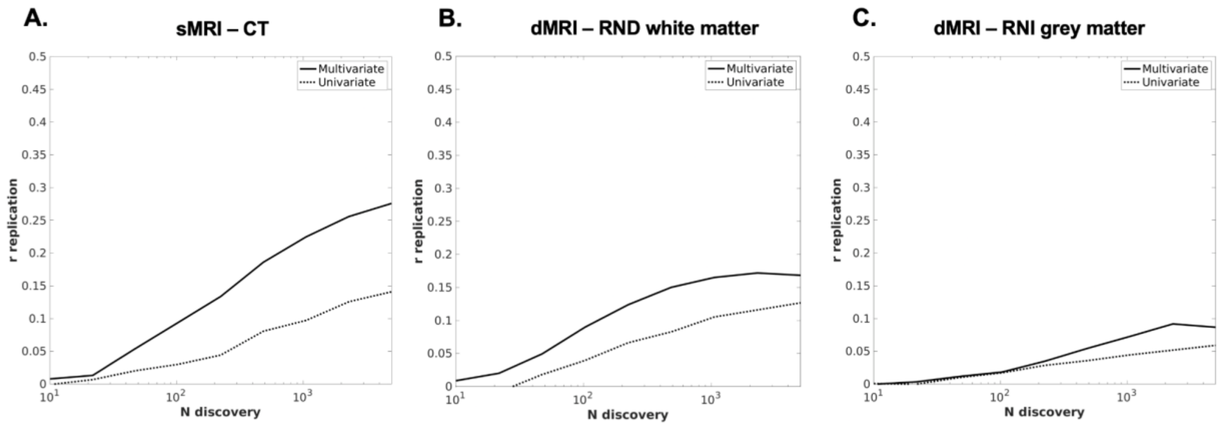


**Supplementary Figure 4.** Replication curves showing out-of-sample prediction performance of general cognition as a function of discovery sample size for the remaining imaging measures not shown in Figure 4 of the main manuscript. The *x*-axis represents discovery sample size in log-scale units and *y*-axis reflects out-of-sample correlation. Multivariate metrics are compared to univariate *r*-values, reflecting the absolute maximum correlation value.

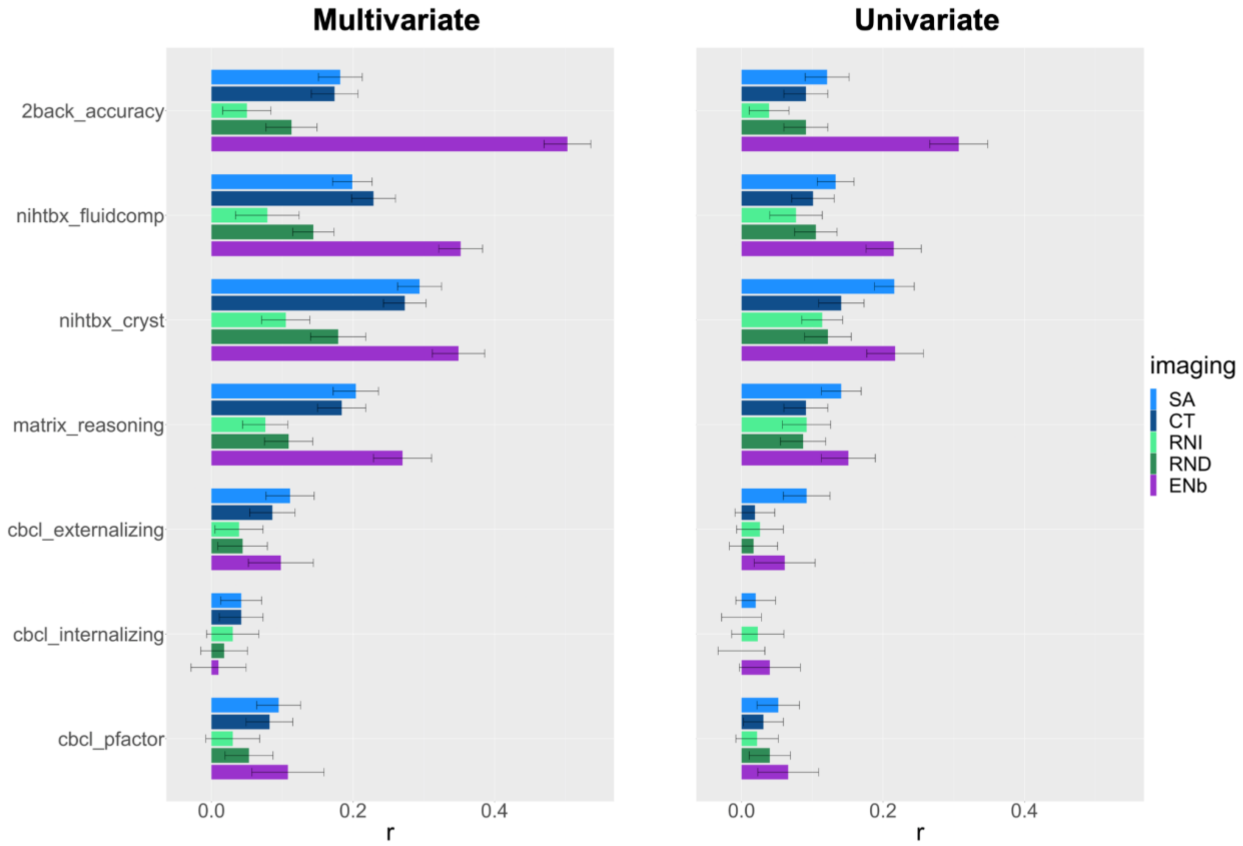


**Supplementary Figure 5**. Out-of-sample prediction performance of 7 other behavioral measures (1 in-scanner cognitive: accuracy on 2-back trials during EN-back task; 3 out-of-scanner cognitive: crystallized and fluid composite scores from the NIH toolbox, matrix reasoning; 3 psychopathology: internalizing/externalizing symptoms and a *p* factor (8) drawn from the Child Behavior Checklist). Error bars reflect standard deviation, adjusted for the 10% sample overlap in test datasets. Abbreviations: SA. surface area; CT. cortical thickness; RND. Restricted directional diffusion within superficial white matter; RNI. Restricted isotropic diffusion intracortically; ENb. Emotional N-back, reflecting the 2- vs. 0-back contrast; CBCL. Child Behavior Checklist. nihtbx_fluidcomp: fluid composite score from NIH toolbox; nihtbx_cryst: crystallized composite score from NIH toolbox.

| Variable | | sMRI | dMRI | tfMRI |
|---|---|---|---|---|
| N | | 11,174 | 10,200 | 5673 |
| N, Female (%) | | 5359 (47.96) | 4941 (48.44) | 2796 (49.29) |
| Mean age in months (sd) | | 119.04 (7.50) | 119.11 (7.50) | 119.8 (7.96) |
| Mean Total Composite Score (sd) | | 86.34 (9.10) | 86.58 (8.99) | 88.75 (7.96) |
| Highest parental education, N (%) | < HS Diploma | 541 (4.84) | 477 (4.68) | 175 (3.08) |
| | HS Diploma/GED | 1050 (9.4) | 922 (9.04) | 349 (6.15) |
| | Some College | 2893 (25.89) | 2617 (25.66) | 1297 (22.86) |
| | Bachelor | 2853 (25.53) | 2620 (25.69) | 1588 (27.99) |
| | Post Grad | 3824 (34.22) | 3552 (34.82) | 2264 (39.91) |
| | NA | 13 (0.12) | 12 (0.12) | 0 (0) |
| Household income N (%) | <50K | 2998 (26.83) | 2667 (26.15) | 1203 (21.21) |
| | ≥ 50K & < 100K | 2897 (25.93) | 2626 (25.75) | 1523 (26.85) |
| | ≥100K | 4334 (38.79) | 4047 (36.98) | 2574 (45.37) |
| | NA | 945 (8.46) | 860 (8.43) | 373 (6.58) |
| Race.4level N (%) | Asian | 259 (2.32) | 225 (2.21) | 149 (2.63) |
| | Black | 1708 (15.29) | 1518 (14.88) | 558 (9.84) |
| | Other/Mixed | 1909 (17.08) | 1722 (16.88) | 911 (16.06) |

| | | | | |
|---|---|---|---|---|
| | White | 7137 (63.87) | 6593 (64.64) | 3999 (70.49) |
| | NA | 161 (1.44) | 142 (1.39) | 56 (0.99) |
| Hispanic N (%) | No | 8750 (78.31) | 7989 (78.32) | 4613 (81.32) |
| | Yes | 2280 (20.40) | 2075 (20.34) | 1060 (18.68) |
| | NA | 144 (1.29) | 136 (1.33) | 0 (0) |

**Supplementary Table 1**. Descriptives of sociodemographic variables for image modality-specific baseline samples when predicting NIH toolbox total composite score.

| predicted variable manuscript | data dictionary variable | ABCD instrument |
|---|---|---|
| nihtbx_totalcomp | nihtbx_totalcomp_uncorrected | abcd_tbss01 |
| 2back_accuracy | tfmri_nb_all_beh_c2b_rate | abcd_mrinback02 |
| nihtbx_fluidcomp | nihtbx_fluidcomp_uncorrected | abcd_tbss01 |
| nihtbx_cryst | nihtbx_cryst_uncorrected | abcd_tbss01 |
| matrix_reasoning | pea_wiscv_trs | abcd_ps01 |
| cbcl_internalizing | cbcl_scr_syn_internal_r | abcd_cbcls01 |
| cbcl_externalizing | cbcl_scr_syn_external_r | abcd_cbcls01 |
| cbcl_pfactor | PF10_lavaan | calculated based on Clark et al (8) |

**Supplementary Table 2.** Behavioral variables used as predictors.

| MRI measure | predicted variable manuscript | emp *k* ratio | mvar *r* | mvar *sd* | uvar *r* | uvar *sd* | *n* full sample | *n* mvar power | *n* uvar power |
|---|---|---|---|---|---|---|---|---|---|
| SA | nihtbx_totalcomp | 0.228 | 0.276 | 0.027 | 0.205 | 0.03 | 11174 | 101 | 185 |
| SA | 2back_accuracy | 0.228 | 0.182 | 0.031 | 0.121 | 0.031 | 11078 | 235 | 534 |
| SA | nihtbx_fluidcomp | 0.228 | 0.199 | 0.028 | 0.133 | 0.026 | 11176 | 196 | 441 |
| SA | nihtbx_cryst | 0.228 | 0.294 | 0.031 | 0.216 | 0.028 | 11230 | 89 | 166 |
| SA | matrix_reasoning | 0.139 | 0.204 | 0.032 | 0.141 | 0.028 | 11168 | 186 | 393 |
| SA | cbcl_internalizing | 0.019 | 0.042 | 0.029 | 0.02 | 0.028 | 11395 | 4447 | >10,000 |
| SA | cbcl_externalizing | 0.019 | 0.111 | 0.034 | 0.092 | 0.033 | 11395 | 635 | 925 |
| SA | cbcl_pfactor | 0.032 | 0.095 | 0.031 | 0.052 | 0.03 | 11394 | 867 | 2900 |
| CT | nihtbx_totalcomp | 0.373 | 0.284 | 0.031 | 0.139 | 0.033 | 11174 | 95 | 404 |
| CT | 2back_accuracy | 0.228 | 0.174 | 0.033 | 0.091 | 0.031 | 11078 | 257 | 946 |
| CT | nihtbx_fluidcomp | 0.228 | 0.229 | 0.031 | 0.101 | 0.03 | 11176 | 147 | 767 |
| CT | nihtbx_cryst | 0.228 | 0.273 | 0.030 | 0.141 | 0.032 | 11230 | 103 | 393 |
| CT | matrix_reasoning | 0.228 | 0.184 | 0.034 | 0.091 | 0.031 | 11168 | 230 | 946 |
| CT | cbcl_internalizing | 0.052 | 0.042 | 0.031 | 0 | 0.028 | 11395 | 4447 | >10,000 |
| CT | cbcl_externalizing | 0.032 | 0.086 | 0.032 | 0.019 | 0.028 | 11395 | 1059 | >10,000 |
| CT | cbcl_pfactor | 0.019 | 0.082 | 0.033 | 0.031 | 0.028 | 11394 | 1165 | 8165 |
| RNI | nihtbx_totalcomp | 0.228 | 0.093 | 0.042 | 0.083 | 0.036 | 10200 | 905 | 1137 |
| RNI | 2back_accuracy | 0.139 | 0.05 | 0.034 | 0.039 | 0.028 | 10166 | 3137 | 5158 |
| RNI | nihtbx_fluidcomp | 0.228 | 0.079 | 0.045 | 0.077 | 0.037 | 10202 | 1255 | 1322 |
| RNI | nihtbx_cryst | 0.228 | 0.105 | 0.034 | 0.114 | 0.029 | 10247 | 710 | 602 |
| RNI | matrix_reasoning | 0.228 | 0.076 | 0.032 | 0.092 | 0.034 | 10202 | 1357 | 925 |
| RNI | cbcl_internalizing | 0.019 | 0.03 | 0.037 | 0.023 | 0.037 | 10400 | 8710 | >10,000 |
| RNI | cbcl_externalizing | 0.139 | 0.039 | 0.034 | 0.026 | 0.033 | 10400 | 5158 | >10,000 |

| | | | | | | | | | |
|---|---|---|---|---|---|---|---|---|---|
| RNI | cbcl_pfactor | 0.139 | 0.03 | 0.038 | 0.022 | 0.03 | 10399 | 8719 | >10,000 |
| RND | nihtbx_totalcomp | 0.228 | 0.163 | 0.039 | 0.127 | 0.034 | 10200 | 293 | 484 |
| FA | nihtbx_totalcomp | 0.228 | 0.157 | 0.044 | 0.133 | 0.036 | 10200 | 316 | 441 |
| RND | 2back_accuracy | 0.228 | 0.113 | 0.036 | 0.091 | 0.031 | 10166 | 612 | 946 |
| RND | nihtbx_fluidcomp | 0.228 | 0.144 | 0.029 | 0.105 | 0.03 | 10202 | 376 | 710 |
| RND | nihtbx_cryst | 0.228 | 0.179 | 0.039 | 0.122 | 0.033 | 10247 | 243 | 525 |
| RND | matrix_reasoning | 0.228 | 0.109 | 0.034 | 0.087 | 0.032 | 10202 | 658 | 1035 |
| RND | cbcl_internalizing | 0.085 | 0.018 | 0.033 | 0 | 0.033 | 10400 | >10,000 | >10,000 |
| RND | cbcl_externalizing | 0.139 | 0.044 | 0.035 | 0.017 | 0.034 | 10400 | 4052 | >10,000 |
| RND | cbcl_pfactor | 0.139 | 0.053 | 0.034 | 0.04 | 0.029 | 10399 | 2792 | 4903 |
| ENb | nihtbx_totalcomp | 0.012 | 0.425 | 0.036 | 0.242 | 0.04 | 5675 | 41 | 132 |
| ENb | 2back_accuracy | 0.019 | 0.503 | 0.033 | 0.307 | 0.041 | 5672 | 29 | 81 |
| ENb | nihtbx_fluidcomp | 0.012 | 0.352 | 0.031 | 0.215 | 0.039 | 5672 | 61 | 168 |
| ENb | nihtbx_cryst | 0.012 | 0.349 | 0.037 | 0.217 | 0.04 | 5672 | 62 | 164 |
| ENb | matrix_reasoning | 0.012 | 0.27 | 0.041 | 0.151 | 0.038 | 5572 | 105 | 342 |
| ENb | cbcl_internalizing | 0.019 | 0.01 | 0.039 | 0.04 | 0.043 | 5670 | >10,000 | 4903 |
| ENb | cbcl_externalizing | 0.004 | 0.098 | 0.046 | 0.061 | 0.043 | 5670 | 815 | 2107 |
| ENb | cbcl_pfactor | 0.004 | 0.108 | 0.051 | 0.066 | 0.043 | 5670 | 671 | 1800 |

**Supplementary Table 3**. Prediction performance per imaging measure and behavioral measure, including NIH toolbox total composite score (presented in manuscript) and seven additional behavioral measures. "Emp *k* ratio" reflects the empirically determined *k* ratio (proportion of principal components used in multivariate prediction of behavior) that optimally predicted behavior in the first 100 iterations of prediction, and was used in the subsequent 100 iterations. '*N* full sample' is the total sample size used in both univariate and multivariate analyses for a given imaging and behavioral variable pair. '*N* mvar/uvar power' columns refer to the sample size required to detect the measured multivariate and univariate effects with 80% power, respectively. Abbreviations: mvar, multivariate. uvar, univariate. SA. surface area; CT. cortical thickness; RND. Restricted directional diffusion within superficial white matter; RNI. Restricted isotropic diffusion intracortically; ENb. Emotional N-back, reflecting the 2- vs. 0-back contrast; CBCL. Child Behavior Checklist. nihtbx_fluidcomp: fluid composite score from NIH toolbox; nihtbx_cryst: crystallized composite score from NIH toolbox.

| Criteria | Instrument | Element value |
|---|---|---|
| | *sMRI* | |
| T1 series passed rawQC | mriqcrp103 | iqc_t1_ok_ser > 0 |
| FreeSurfer QC not failed | abcd_fsurfqc01 | fsqc_qc ~= 0 |
| Derived results exist | abcd_smrip202 | smri_t1w_scs_cbwmatterlh ~= NA |
| | *dMRI* | |
| dMRI series passed rawQC | mriqcrp103 | iqc_dmri_ok_ser > 0 |
| dMRI total number of repetitions | mriqcrp103 | iqc_dmri_ok_nreps >= 103 OR (mri_info_manufacturer = Philips AND iqc_dmri_ok_ser >= 2 AND iqc_dmri_ok_nreps = 51) |
| T1 series passed rawQC | mriqcrp103 | iqc_t1_ok_ser > 0 |
| dMRI B0 unwarp available | abcd_auto_postqc01 | apqc_dmri_bounwarp_flag == 1 |
| FreeSurfer QC not failed | abcd_fsurfqc01 | fsqc_qc ~= 0 |
| dMRI manual post-processing QC not failed | abcd_dmriqc01 | dmri_postqc_qc ~= 0 |
| dMRI registration to T1w | abcd_auto_postqc01 | apqc_dmri_regt1_rigid < 17 |
| dMRI dorsal cutoff score | abcd_auto_postqc01 | apqc_dmri_fov_cutoff_dorsal < 47 |
| dMRI ventral cutoff score | abcd_auto_postqc01 | apqc_dmri_fov_cutoff_ventral < 54 |
| Derived results exist | abcd_drsip201 | dmri_rsirnd_fib_allfib ~= NA |

| *tfMRI - EN back* | | |
|---|---|---|
| nBack tfMRI series passed rawQC | mriqcrp103 | iqc_nback_ok_ser > 0 |
| T1 series passed rawQC | mriqcrp103 | iqc_t1_ok_ser > 0 |
| nBack behavior passed | abcd_nback02 | tfmri_nback_beh_performflag == 1 |
| nBack degrees of freedom > 200 | nback_bwroi02 | tfmri_nback_all_b_dof > 200 |
| nBack E-prime timing match OR ignore E-prime mismatch | mriqcrp302 | iqc_nback_ep_t_series_match == 1 \|\| eprime_mismatch_ok_nback == 1 |
| fMRI B0 unwarp available | abcd_auto_postqc01 | apqc_fmri_bounwarp_flag == 1 |
| FreeSurfer QC not failed | abcd_fsurfqc01 | fsqc_qc ~= 0 |
| fMRI manual post-processing QC not failed | abcd_fmriqc01 | fmri_postqc_qc ~= 0 |
| fMRI registration to T1w | abcd_auto_postqc01 | apqc_fmri_regt1_rigid < 19 |
| fMRI dorsal cutoff score | abcd_auto_postqc01 | apqc_fmri_fov_cutoff_dorsal < 65 |
| fMRI ventral cutoff score | abcd_auto_postqc01 | apqc_fmri_fov_cutoff_ventral < 60 |
| Derived results exist | nback_bwroi02 | tfmri_nback_all_4 ~= NA |

**Supplementary Table 4**. Inclusion/exclusion criteria for image include flags per modality.

### SUPPLEMENTARY REFERENCES


1. H. Garavan, *et al.*, Recruiting the ABCD sample: Design considerations and procedures. *Dev. Cogn. Neurosci.* **32**, 16–22 (2018).

2. N. D. Volkow, *et al.*, The conception of the ABCD study: From substance use to a broad NIH collaboration. *Dev. Cogn. Neurosci.* **32**, 4–7 (2018).

3. M. Luciana, *et al.*, Adolescent neurocognitive development and impacts of substance use: Overview of the adolescent brain cognitive development (ABCD) baseline neurocognition battery. *Dev. Cogn. Neurosci.* **32**, 67–79 (2018).

4. N. Akshoomoff, *et al.*, VIII. NIH Toolbox Cognition Battery (CB): composite scores of crystallized, fluid, and overall cognition. *Monogr. Soc. Res. Child Dev.* **78**, 119–132 (2013).

5. R. K. Heaton, *et al.*, Reliability and validity of composite scores from the NIH Toolbox Cognition Battery in adults. *J. Int. Neuropsychol. Soc.* **20**, 588–598 (2014).

6. S. Marek, *et al.*, Reproducible brain-wide association studies require thousands of individuals. *Nature* **603**, 654–660 (2022).

7. D. Wechsler, *Wechsler Intelligence Scale for Children*, E. Pearson 5th, Ed. (Bloomington, MN, 2014).

8. D. A. Clark, *et al.*, The General Factor of Psychopathology in the Adolescent Brain Cognitive Development (ABCD) Study: A Comparison of Alternative Modeling Approaches. *Clin. Psychol. Sci.* **9**, 169–182 (2021).

9. B. J. Casey, *et al.*, The Adolescent Brain Cognitive Development (ABCD) study: Imaging acquisition across 21 sites. *Dev. Cogn. Neurosci.* **32**, 43–54 (2018).

10. D. J. Hagler Jr, *et al.*, Image processing and analysis methods for the Adolescent Brain Cognitive Development Study. *Neuroimage* **202**, 116091 (2019).

11. A. M. Dale, B. Fischl, M. I. Sereno, Cortical surface-based analysis. I. Segmentation and surface reconstruction. *Neuroimage* **9**, 179–194 (1999).

12. B. Fischl, M. I. Sereno, A. M. Dale, Cortical surface-based analysis. II: Inflation, flattening, and a surface-based coordinate system. *Neuroimage* **9**, 195–207 (1999).

13. B. Fischl, *et al.*, Automatically parcellating the human cerebral cortex. *Cereb. Cortex* **14**, 11–22 (2004).

14. B. Fischl, A. M. Dale, Measuring the thickness of the human cerebral cortex from magnetic resonance images. *Proc. Natl. Acad. Sci. U. S. A.* **97**, 11050–11055 (2000).

15. J. Jovicich, *et al.*, Reliability in multi-site structural MRI studies: effects of gradient non-linearity correction on phantom and human data. *Neuroimage* **30**, 436–443 (2006).

16. N. S. White, T. B. Leergaard, H. D'Arceuil, J. G. Bjaalie, A. M. Dale, Probing tissue microstructure with restriction spectrum imaging: Histological and theoretical validation. *Hum. Brain Mapp.* **34**, 327–346 (2013).

17. N. S. White, *et al.*, Diffusion-weighted imaging in cancer: physical foundations and applications of restriction spectrum imaging. *Cancer Res.* **74**, 4638–4652 (2014).

18. C. E. Palmer, *et al.*, Microstructural development from 9 to 14 years: Evidence from the ABCD Study. *Dev. Cogn. Neurosci.* **53**, 101044 (2022).

19. P. J. Basser, C. Pierpaoli, Microstructural and physiological features of tissues elucidated by quantitative-diffusion-tensor MRI. *J. Magn. Reson.* **213**, 560–570 (1996).

20. P. J. Basser, J. Mattiello, D. LeBihan, Estimation of the effective self-diffusion tensor from the NMR spin echo. *J. Magn. Reson. B* **103**, 247–254 (1994).

21. A. O. Cohen, *et al.*, When Is an Adolescent an Adult? Assessing Cognitive Control in Emotional and Nonemotional Contexts. *Psychol. Sci.* **27**, 549–562 (2016).

22. A. O. Cohen, M. I. Conley, D. V. Dellarco, B. J. Casey, The impact of emotional cues on short-term and long-term memory during adolescence. *Proceedings of the Society for Neuroscience. San Diego* (2016).

23. W. Zhao, *et al.*, Task fMRI paradigms may capture more behaviorally relevant information than resting-state functional connectivity. *Neuroimage* **270**, 119946 (2023).

24. D. A. Fair, *et al.*, Correction of respiratory artifacts in MRI head motion estimates. *Neuroimage* **208**, 116400 (2020).

25. J. S. Siegel, *et al.*, Statistical improvements in functional magnetic resonance imaging analyses produced by censoring high-motion data points. *Hum. Brain Mapp.* **35**, 1981–1996 (2014).

26. C. Sripada, *et al.*, Prediction of neurocognition in youth from resting state fMRI. *Mol. Psychiatry* **25**, 3413–3421 (2020).

27. T. Spisak, U. Bingel, T. D. Wager, Multivariate BWAS can be replicable with moderate sample sizes. *Nature* **615**, E4–E7 (2023).

28. W. H. Bondy, W. Zlot, The Standard Error of the Mean and the Difference between Means for Finite Populations. *Am. Stat.* **30**, 96–97 (1976).

29. S. B. Hulley, S. R. Cummings, W. S. Browner, D. G. Grady, T. B. Newman, *Designing Clinical Research* (Lippincott Williams & Wilkins, 2013).
